# Effects of an IgE receptor polymorphism acting on immunity, susceptibility to infection and reproduction in a wild rodent

**DOI:** 10.1101/841825

**Authors:** Klara M Wanelik, Mike Begon, Janette E Bradley, Ida M Friberg, Joseph A Jackson, Christopher H Taylor, Steve Paterson

## Abstract

The genotype of an individual is an important predictor of their immune function, and subsequently, their ability to control or avoid infection and ultimately contribute offspring to the next generation. However, the same genotype, subjected to different intrinsic and/or extrinsic environments can also result in different phenotypic outcomes, which can be missed in controlled laboratory studies. Natural wildlife populations, which capture both genotypic and environmental variability, provide an opportunity to more fully understand the phenotypic expression of genetic variation. We identified a synonymous polymorphism in the high-affinity Immunoglobulin E (IgE) receptor (GC and non-GC haplotypes) that has sex-dependent effects on immune gene expression, susceptibility to infection and reproductive success of individuals in a natural population of field voles (*Microtus agrestis*). We found that the effect of the GC haplotype on the expression of immune genes differed between sexes. While males with the GC haplotype had upregulated more pro-inflammatory genes, including the cytokine *Il33*, females had upregulated more anti-inflammatory genes, including the cytokine inhibitor *Socs3*. Furthermore we found an effect of the GC haplotype on the probability of infection with a common microparasite*, Babesia microti*, in females - with females carrying the GC haplotype being more likely to be infected. Finally, we found an effect of the GC haplotype on reproductive success in males - with males carrying the GC haplotype having a lower reproductive success. This is a rare example of a polymorphism whose consequences we are able to follow across immunity, infection and reproduction for both males and females in a natural population.

## Introduction

In order for an individual to control or avoid infection and ultimately contribute offspring to the next generation, they must have a well-functioning immune system [1]. An individual’s immune function is in part determined by their genotype (e.g. [2–4]). Individuals within a population differ from each other in the genotypes they carry, with the potential for these different genotypes to affect the activity of different immune components, and thereby influence susceptibility to infection and reproduction and survivorship. Indeed, variability in immune responses to infection is well-characterised. The phenotypic expression of a genotype is dependent on the environment, including the sex of an individual (intrinsic environment) and exposure to infection (extrinsic environment). Since capturing genotypic and environmental variability in controlled laboratory studies is problematic, natural wildlife populations have been proposed as models in which to understand the phenotypic expression of genetic variation and its consequences for susceptibility to infection and reproduction.

Studies of natural populations have explored the effects of genotype on immune phenotype, and have observed consequences for susceptibility to infection. Most notably, variability in the genes of the Major Histocompatibility Complex (MHC) has been associated with resistance to intestinal nematodes in domestic sheep [2] and with resistance to malaria, hepatitis, and AIDS in humans [5–7]. The role of variability elsewhere in the genome, for shaping immune phenotype, has also been studied [3,4]. However, it remains challenging to follow the consequences of a genotypic effect for immunity, infection and reproduction and to account for any sex-dependent expression of a genotype. This is because of the difficulty in obtaining phenotypic data across immune, infection and reproductive traits, especially for large enough sample sizes to test for data-hungry genotype by sex interactions [8]. In many cases, sex, and other environmental factors are considered as a confounding variable to be controlled for in order not to hide any subtle genetic associations [2]. Other studies focus on a single sex for the sake of simplicity [9]. More recently, however, there has been a growing body of large-scale field studies of natural populations able to apply genetic and immunological methods to follow large numbers of individuals, exposed to a challenging environment and with varying genetic backgrounds, throughout their lives. This allows us to make a more complete assessment of the impacts of genotype throughout the life of an individual, whether male or female. For example, Graham et al. 2010 [10] found evidence for heritable variation in immunity associated with sex-dependent effects on Soay sheep reproduction. However, we know of no documented example of a polymorphism affecting immunity, susceptibility to infection and reproduction in a natural population investigated in males and females. Here, we use a wild rodent, the field vole (*Microtus agrestis*), as a model in which to do this. Wild rodents, in particular, offer an opportunity to quickly follow large numbers of individuals throughout their lives, given their short lifespans. They also offer the opportunity to draw on the immunological and genetics resources developed for laboratory rodents, while providing a much more realistic ecological model of human populations [11,12].

Immunoglobulin E (IgE) mediated responses are associated with defence against helminths [13] and with allergy [14]. They are controlled by the high-affinity IgE receptor, FCER1, which is found on the surface of various immune effector cells e.g. mast cells, basophils and eosinophils [15]. In humans, naturally occurring polymorphisms in FCER1 are known to affect an individual’s serum IgE levels, with consequences for their susceptibility to infection [16,17] and their risk of developing inflammatory disease [18–21]. Furthermore, sex differences in serum IgE levels [22] and the incidence of IgE-mediated inflammatory disease [23] have been documented in humans, suggesting that any polymorphism in this pathway is likely to experience different contexts in males and females. Indeed, one study found evidence for a polymorphism in the *Fcer1a* gene (the alpha chain of FCER1) whose association with susceptibility to systematic lupus erythematosus (a chronic inflammatory disease) differed between males and females [24].

In a previous study of a natural population of *M. agrestis*, we found that males carrying the GC haplotype of the *Fcer1a* gene expressed the transcription factor GATA3 at a lower level than males carrying non-GC haplotypes [3]. GATA3 is a biomarker of tolerance to macroparasites in mature males in our population (macroparasite infection gives rise to increased expression of GATA3, which gives rise to improved body condition and survival [9]). Here, we explore the effects of this GC haplotype further in both males and females to analyse the effect of genotype acting across immune expression, infection susceptibility and reproductive success.

## Results

We sampled a natural population of *M. agrestis* in Kielder Forest, Northumberland, United Kingdom, over three years (2015-2017) and across seven different sites. Our study involved a cross-sectional component (*n* = 317 destructively sampled voles) and a longitudinal component (*n* = 850 marked individuals monitored through time, with *n* = 2,387 sampling points). We tested the consequences of the GC haplotype of the *Fcer1a* gene using both cross-sectional and longitudinal components of our study. As well as the GC haplotype (present at a frequency of 0.08), three other haplotypes were present in our study population: AC haplotype, AT haplotype and GT haplotype, present at frequencies of 0.81, 0.10 and 0.01 respectively. The two SNPs composing the haplotype were found to be tightly linked (r^2^ = 0.50; D’ = 0.70).

### The GC haplotype has effects on inflammation that differ between sexes

In humans, naturally occurring polymorphisms in the *Fcer1a* gene have previously been linked to inflammatory disease [18–21]. Therefore, we used the cross-sectional component of our study to test the effects of the GC haplotype on inflammation in males and females. Differential gene expression (DGE) analysis performed on unstimulated splenocytes taken from 53 males and 31 females assayed by RNASeq, showed that the identity of top-responding genes differed between the sexes. In males, the top-responding immune gene was the cytokine, *Il33* (log fold change (logFC) = 2.76, *p* = 0.00000015, *q* = 0.00048; Appendix 1—table 2) while in females it was the suppressor of cytokine signalling *Socs3* (logFC = 1.07, *p* = 0.000061, *q* = 0.05; Appendix 1—table 3). Looking at the ranking of each top-responding gene in the opposite sex strengthens the case for differing effects in males and females, with *Il33* ranked markedly lower in females (rank = 8224/12904, logFC = 0.29, *p* = 0.70, *q* = 1.00) and *Socs3* markedly lower in males (rank = 10886/12904, logFC = −0.05, *p* = 0.84, *q* = 1.00). *Il33* is commonly associated with the anti-helminthic response [25] and *Socs3* with regulation of the immune response more broadly [26]. Given the link between *Il33* and the antihelminthic response (and more generally, IgE-mediated responses and the antihelminthic response), we repeated the DGE analysis while controlling for cestode burden, but this had little effect on our results (same top-responding immune genes; see Appendix 1—table 4 & 5), suggesting that these effects were not driven by differences in cestode infection.

Both *Il33* and *Socs3* also share an association with the inflammatory response [26,27]. While *Il33* positively regulates this response (appearing in the gene set GO:0050729), *Socs3* negatively regulates it (GO:0050728). To test whether these effects on the inflammatory response were limited to these genes or were more widespread, we performed a gene set enrichment analysis (GSEA) which looked at the rankings of all genes present in each of these gene sets (GO:0050729 and GO:0050728). This analysis showed that both males and females upregulated their expression of both gene sets. However, males showed a stronger signal for genes which positively regulate the inflammatory response (GO:0050729: *p* = 0.007; GO:0050728: *p* = 0.04; Figure 1A, upper panel) while females showed a stronger signal for genes which negatively regulate the inflammatory response (GO:0050728: *p* = 0.001; GO:0050728: *p* = 0.04; Figure 1A, lower panel).

**Figure 1.**
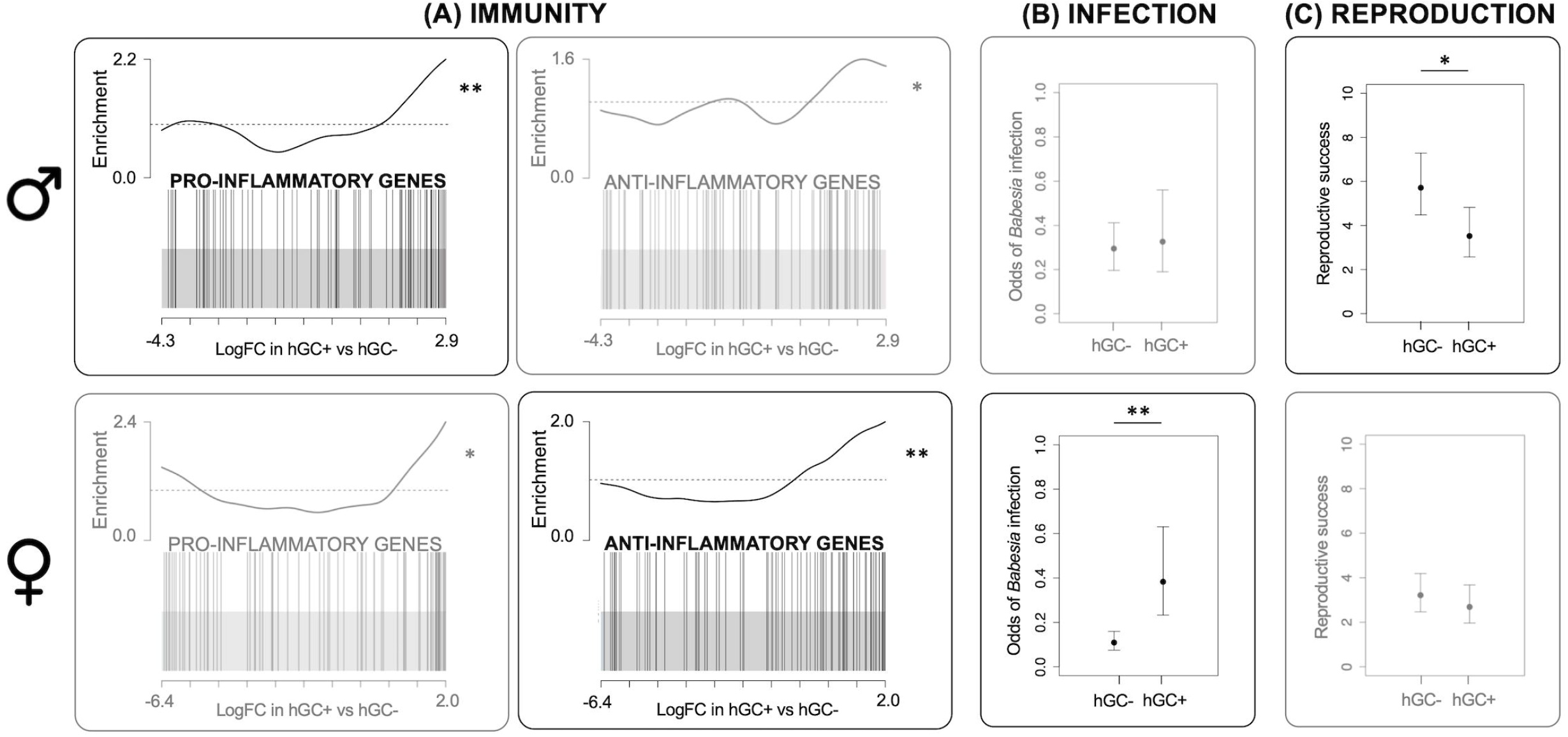
Effects of GC haplotype (hGC). Upper panel: males. Lower panel: females: (A) Unstimulated immune gene expression: Barcode plots showing enrichment of the GO terms GO:0050729 (pro-inflammatory genes) and GO:0050728 (anti-inflammatory genes) in unstimulated splenocytes taken from individuals with (hGC+) vs. without (hGC-) the haplotype, showing that males with the haplotype have a pro-inflammatory bias, whereas females have an anti-inflammatory bias. In each plot, x-axis shows log fold change (logFC) in hGC+ vs. hGC-, black bars represent genes annotated with the GO terms and the worm shows relative enrichment. (B) Susceptibility to infection: Association between hGC and the odds of infection with *Babesia microti*, showing that females with the haplotype have an increased susceptibility to infection. (C) Reproduction: Association between hGC and reproductive success, showing that males with the haplotype have lower reproductive success (error bars represent ± standard error; * indicates *p* < 0.05; ** indicates *p* < 0.01).

To further explore these effects on the inflammatory response, we used an independent dataset for males and females whose spleens were stimulated with two immune agonists and assayed by Q-PCR (for a panel of 18 immune genes with limited redundancy; see Methods for how these genes were selected). From this *ex vivo* assay, one can gain insight into the types of immune response that could be made to a pathogen *in vivo* (see Methods for details). Although our panel of genes did not include *Il33* or *Socs3*, it did include other genes associated with the inflammatory response including *Il17a, Ifng, Il1b, Il6* and *Tnfa*. We found that the effect of the GC haplotype on the stimulated expression levels of *Il17a*, a pro-inflammatory cytokine involved in the antibacterial response, displayed a significant interaction with sex (genotype by sex interaction in follow-up LMM, LRT *p* = 0.001). While females with the GC haplotype expressed lower levels of *Il17a* (estimate = −0.12, 95% CI = −0.19 – −0.04), males with the GC haplotype did not differ in their expressed levels of *Il17a* (estimate = 0.03, 95% CI includes zero = −0.02 – 0.08; Appendix 1—table 6 & 7).

### The GC haplotype affects oxidative stress

Inflammation and oxidative stress are closely related and often lead to pathology [28]. To test whether the effects of the GC haplotype on inflammation may be having knock-on effects on oxidative stress, we again used the cross-sectional component of our study. We tested the effect of the GC haplotype on antioxidant superoxide dismutase 1 (SOD1) enzymatic activity in males and females. SOD1 is one of the main antioxidative enzymes within the antioxidative enzyme system [29]. We found that (irrespective of sex) individuals with the AT haplotype had lower SOD1 activity when compared with other haplotypes, including the GC haplotype (*p* = 0.04; Appendix 1—table 8).

### The GC haplotype affects susceptibility to *Babesia* infection in females

We then used the longitudinal component of our study to test the effects of the GC haplotype on the probability of infection with common parasites known to infect our study population: two micro-parasites *Babesia microti* and *Bartonella* spp. and a range of macroparasites (ticks, fleas and cestodes). We found an effect of the GC haplotype on the probability of infection with *B. microti* which displayed a marginally significant interaction with sex (genotype by sex interaction, LRT *p* = 0.048; Appendix 1—table 9). While females with the GC haplotype were more likely to be infected with *B. microti* (log odds ratio = 1.25, 95% CI = 0.44 – 2.07; Figure 1B, lower panel), males with the GC haplotype did not differ in their susceptibility to infection (log odds ratio = 0.14, 95% CI includes zero = −0.76 – 1.03; Figure 1B, upper panel). However, we found no effect of the haplotype (interactive or not) on the probability of infection with the other parasites in our population (Appendix 1—table 11 & 17).

### The GC haplotype reduces male reproductive success

We genotyped both cross-sectional and longitudinal voles for 114 SNPs and used this dataset to construct a pedigree. We estimated each individuals’ reproductive success by counting the number offspring it produced according to this pedigree. We then tested the effects of the GC haplotype on this measure of reproductive success. We tried to run a single model with both sexes (as above) but resulting residual variances differed significantly between the sexes, which reflects the fact that male and female reproduction are inherently different traits. This made it impossible to formally test for a genotype by sex interaction and so instead we ran a separate model for each sex. We found that males with the GC haplotype had an average of 2.2 fewer offspring than males without the haplotype (*p* = 0.04; Appendix 1—table 12; Figure 1C, upper panel). Despite a larger sample size and lower variability in reproductive success, females with the GC haplotype did not differ significantly in their number of offspring (genotype term did not appear in best model; Appendix 1—table 13; Figure 1C, lower panel). This is suggestive of a significant genotype by sex interaction. We ran the same models including microparasite variables, but this had little effect on our results (see Appendix 1—table 14 & 15), suggesting that these effects were not driven by differences in microparasite infection.

## Discussion

In this study, we describe a polymorphism in the high-affinity receptor for IgE (*Fcer1a*) with effects that act across immune gene expression, oxidative stress, susceptibility to infection and ultimately reproductive success in a natural population. This begins to reveal how genotypic effects can have multiple effects on different phenotypic or life-history traits for organisms living in the natural environment. Interestingly, effects often differed between sexes, with evidence for opposing effects in the sexes (unstimulated immune gene expression), and an effect present in one sex but not the other (stimulated immune gene expression, susceptibility to infection and reproductive success). However, we cannot rule out that, in the case of susceptibility to infection and reproductive success, there was a smaller effect present in the opposite sex (in the same or opposite direction) which our analysis did not have sufficient power to detect.

In humans, naturally occurring polymorphisms in high-affinity IgE receptor have previously been linked to inflammatory disease [18–21]. Our work is consistent with this and suggests differing effects of this polymorphism on the inflammatory phenotype of males and females. While males with the GC haplotype showed a pro-inflammatory bias, females with the haplotype showed an anti-inflammatory bias. A previous study on another polymorphism in the human ortholog of *Fcer1a* also found evidence for sex-dependent effects on inflammatory disease [24]. In order to fully understand this genotype by sex interaction, it is important to consider background differences in the inflammatory phenotypes of males and females. Females in this study were pregnant for a large proportion of their lives (57% of post-juvenile females were pregnant or lactating at the time of capture in the longitudinal component of our study). In mammals, including humans, pregnancy has been shown to be a largely anti-inflammatory state, with pregnant females suppressing inflammation. This is to protect the foetus from attack by the mother’s immune system [30]. While testosterone in males has an anti-inflammatory effect [31], it also drives males to use more space [32] and to be more aggressive [33]. These behaviours might increase their rates of contact with other individuals, and with their environment, and hence their likelihood of infection and subsequent immune stimulation. We suggest that the polymorphism may be exaggerating background sex differences in inflammatory activity. Sex hormones have been implicated in the differential expression of some autosomal genes in males and females [34], and could be driving these opposing effects. Some of the differences in immune phenotype that we observed may also be driven by difference in parasite infection (although we accounted for cestode burden in a follow-up analysis, we cannot rule this out).

The effect of *Fcer1a* polymorphism on inflammatory responses may also, in turn, affect individual susceptibility to infection. This is consistent with a previous association found in this system between IgE-mediated responses and defence against helminths [13]. Although we did not find an association between macroparasite burden (including cestode burden) and the GC haplotype in this study, in our previous study, we showed that males with the GC haplotype had a lower level of an immunological marker of tolerance to ticks, fleas and adult cestodes (i.e., they were less tolerant) [3]. Here, we use data on infection incidence to show that, in a different context, the same haplotype is also associated with resistance to a microparasite, *B. microti*, in females. Although observational studies of this kind are not able to identify causal relationships, a realistic scenario is that females with the GC haplotype are more likely to be infected with *B. microti* (i.e., they are less resistant) as a result of a lower pro-inflammatory to anti-inflammatory cytokine ratio. Pro-inflammatory cytokines (e.g., IL-6, IFN-γ, and TNF-α) may help to resist *B. microti* infection [35]. A lack or imbalance of these cytokines may hamper this resistance. The panel of parasites that we have measured is not exhaustive and previous studies have highlighted the important role of species interactions in the parasite community in driving infection risk in this study population [36]. *B. microti* may therefore represent a community of parasites, to which this haplotype affects resistance in females.

A fitness consequence of the GC haplotype was observed in reduced male reproductive success. A realistic scenario is that the larger cost incurred by males with the GC haplotype due to inflammation reduced their reproductive value, as reproducing and mounting a pro-inflammatory response are both costly activities [37]. Another realistic scenario is that the positive effect of the GC haplotype on levels of oxidative stress (as indicated by a tendency for higher SOD1 activity) may have had a more detrimental effect in males, damaging sperm and reducing fertilising success. Sperm competition can be an important feature in the reproduction of microtines, e.g., meadow vole (*M. pennsylvanicus*) [38], with males having to produce many sperm of high quality in order to successfully outcompete other males. Sperm are also highly susceptible to damage by reactive oxygen species (ROS) [39]. In addition, the heightened level of oxidative stress may have impaired olfactory sexual signals, e.g., excretion of major urinary proteins (MUPs) making males less attractive to females and negatively impacting on mating success [40].

Despite the immune system playing an important role in determining the health and reproductive fitness of an individual, natural selection has not converged on a single immune optimum. Instead, individuals in natural populations vary widely in their response to infection. Understanding why immunoheterogeneity is maintained is a key question in eco-immunology. Antagonistic selection, where a mutation is beneficial in some environmental contexts and harmful in others, is thought to be one mechanism by which balancing selection is generated, and genetic variation in immunity can be maintained within natural populations [10]. In samples collected between 2008-2010, we previously identified four haplotypes at this locus, GC, AC, AT and GT, at frequencies of 0.12, 0.76, 0.07 and 0.05 respectively [3]. Seven years on, these frequencies have remained relatively unchanged (0.08, 0.81, 0.10 and 0.01). The fact that the GC haplotype remains in the population, despite its detrimental effects on male reproductive success and female infection susceptibility, suggests that it may be under balancing selection.

In order to be under balancing selection though, the GC haplotype would need to confer some advantage. We were not able to find evidence for any such advantage in this study, but we can speculate on one. Microtines show cyclical population dynamics, alternating between a “peak” phase (associated with good conditions and high vole densities) and a “crash” phase (poor conditions and low vole densities). Selection pressures acting during these two phases will be very different. We found that GC haplotype was associated with a pro-inflammatory bias in males, indicating higher investment in promoting inflammation. Thus, one possible scenario is that higher investment in promoting inflammation is disadvantageous in the “peak” phase when the risk of parasitism is low due to dilution effects, and it pays not to waste energy mounting a stronger immune response that is unnecessary. However, it might be advantageous in the “crash” phase when conditions are poor, and the risk of parasitism is high due to the time lagged transmission of parasite infectious stages that have accumulated during the peak phase. Vole numbers were extremely low in our study area between 2012 and 2013 (indicating a “crash phase”). In 2014, vole numbers began to increase and by the beginning of 2015 (the start of our study) had reached a peak. Although vole numbers declined throughout our study period, they never reached the extremely low numbers recorded in 2012-2013 (X. Lambin, personal communication, 2016). This suggests that we sampled in the region of a “peak” phase, when this investment was disadvantageous, and that if we sampled during a true “crash” phase we might detect an advantage to the GC haplotype in males.

There is a growing interest in human genomic (or precision) medicine, with the potential to use a patient’s genotypic information to personalise their treatment. What we have shown here demonstrates that considering genotype in isolation can be misleading, as the same polymorphism can have different outcomes not only for the immune gene expression, but the susceptibility to infection and ultimate reproductive success of males and females.

## Materials and Methods

We studied *M. agrestis* in Kielder Forest, Northumberland, UK, using live-trapping of individual animals from natural populations. Trapping was performed from 2015-2017 across seven different sites, each a forest clear-cut. Access to the sites was provided by the Forestry Commission. At each site, 150-197 Ugglan small mammal traps (Grahnab, Gnosjo, Sweden) were laid out in a grid spaced approximately 3-5 m apart. Our study was divided into longitudinal and cross-sectional components.

### Ethics statement

All animal procedures were performed with approval from the University of Liverpool Animal Welfare Committee and under a UK Home Office licence (PPL 70/8210 to S.P.).

### Longitudinal data

Every other week, traps were checked every day, once in the morning and once in the evening. Newly trapped field voles were injected with a Passive Integrated Transponder (PIT) tag (AVID, Uckfield, UK) for unique identification. This approach allowed us to build up a longitudinal record for voles that were caught on multiple occasions. A total of 850 voles were individually marked in this way. We also took a small tissue sample from the tail, for genotyping of individuals and a drop of blood from the tail which we put into 500 μl of RNAlater (Fisher Scientific, Loughborough, UK) for use in parasite detection (see below).

#### Parasite detection

We quantified infections by microparasites (*Babesia microti* and *Bartonella* spp.) in blood samples taken from longitudinal animals using SYBR green based two-step reverse transcription quantitative PCR (Q-PCR) targeted at pathogen ribosomal RNA genes. Expression values were normalized to two host endogenous control genes: *Ywhaz* and *Actb*. Blood samples were derived from tail bleeds. RNA was extracted from blood samples stored in RNAlater at −70°C using the Mouse RiboPure Blood RNA Isolation Kit (ThermoFisher, Waltham, Massachusetts), according to manufacturer’s instructions, and DNAse treated. It was then converted to cDNA using the High-Capacity RNA-to-cDNA™ Kit (ThermoFisher), according to manufacturer’s instructions. *B. microti* primer sequences targeting the 18S ribosomal RNA gene were as follows CTACGTCCCTGCCCTTTGTA (forward primer sequence) and CCACGTTTCTTGGTCCGAAT (reverse primer sequence). *Bartonella* spp. primer sequences targeting the 16S ribosomal RNA gene were as follows GATGAATGTTAGCCGTCGGG (forward primer sequence) and TCCCCAGGCGGAATGTTTAA (reverse primer sequences). Assays were pipetted onto 384 well plates by a robot (Pipetmax; Gilson, Middleton, Wisconsin) using a custom programme and run on a QuantStudio 6-flex Real-Time PCR System (ThermoFisher) at the machine manufacturers default real-time PCR cycling conditions. Reaction size was 10 μl, incorporating 1 μl of template (diluted 1/20) and PrecisionFAST qPCR Master Mix (PrimerDesign, Chandler’s Ford, UK) with low ROX and SYBR green and primers at the machine manufacturer’s recommended concentrations. Alongside the pathogen assays we ran assays for two host genes (see above) that acted as endogenous controls. For the pathogen assays we used as a calibrator sample a pool of DNA extracted from 154 blood samples from different *M. agrestis* at our study sites in 2015 and 2016; these DNA extractions were carried out using the QIAamp UCP DNA Micro Kit (Qiagen, Hilden, Germany) following manufacturer’s instructions. For the *Ywhaz* and *Actb* assays the calibrator sample was the same cDNA calibrator sample as described below for other host gene expression assays. Relative expression values (normalised to the two host endogenous control genes and presumed to relate to the expression of pathogen ribosomal RNA genes and in turn to the force of infection) used in analyses below are RQ values calculated by the QuantStudio 6-flex machine software according to the ΔΔCt method, indexed to the calibrator samples. Melting curves and amplification plots were individually inspected for each well replicate to confirm specific amplification. We validated our diagnostic results by comparing our PCR RQ values to independent data for a subset of voles from the cross-sectional component of our study with mapped genus-level pathogen reads from RNASeq analysis of blood samples (*n* = 44), finding the two data sets strongly corroborated each other (*Bartonella* spp., Spearman’s *ρ* = 0.79, *p* < 0.001, *n* = 44; *B. microti*, Spearman’s *p* = 0.88, *ρ* < 0.001, *n* = 43).

### Cross-sectional data

For the cross-sectional component of the study (which ran from 2015-2016 only), animals were captured and returned to the laboratory where they were killed by a rising concentration of CO_2_, followed by exsanguination. As our UK Home Office licence did not allow us to sample overtly pregnant females in this way, there are fewer females than males present in this dataset. This component of the study allowed us to take a more comprehensive set of measurements, and to culture cells and perform stimulatory assays on them. A small tissue sample was taken from the ear of cross-sectional animals for genotyping (see below).

#### Parasite detection

After culling, the fur of cross-sectional animals was examined thoroughly under a binocular dissecting microscope to check for ectoparasites, which were recorded as direct counts of ticks and fleas. Guts of cross-sectional animals were transferred to 70% ethanol, dissected and examined under a microscope for gastrointestinal parasites. Direct counts of cestodes (by far the dominant endohelminths in biomass) were recorded.

#### SOD1 measurement

Given the well-established link between inflammation, oxidative stress and pathology [28], we measured antioxidant enzymatic activity in blood samples taken from cross-sectional animals. We chose to measure superoxide dismutase 1 (SOD1) because it is one of the main antioxidative enzymes within the antioxidative enzyme system [29], and is clearly linked to changes in immune gene expression in laboratory mice [41]. Assays were carried out using the Cayman Superoxide Dismutase kit and following the manufacturer’s instructions except where otherwise indicated below. Blood pellets from centrifuged cardiac bleeds were stored at −70°C and thawed on ice prior to assay. A 20 μl aliquot from each pellet was lysed in 80 μl of ultrapure water and centrifuged (10000 G at 4°C for 15 minutes) and 40 μl of the supernatant added to a 1.6 × volume of ice-cold chloroform/ethanol (37.5/62.5 (v/v)) (inactivating superoxide dismutase 2). This mixture was then centrifuged (2500 G at 4°C for 10 minutes) and the supernatant removed and immediately diluted 1:200 in kit sample buffer. A seven-point SOD activity standard was prepared (0, 0.005, 0.010, 0.020, 0.030, 0.040, 0.050 U/ml) and the assay conducted on a 96 well microplate with 10 μl of standard or sample, 200 μl of kit radical detector solution and 20 μl of xanthine oxidase (diluted as per manufacturer’s instructions) per well. Plates were then covered with a protective film, incubated at room temperature for 30 mins on an orbital shaker and read at 450 nm on a VERSAmax tuneable absorbance plate reader (Molecular Devices, San Jose, California), subtracting background and fitting a linear relationship to the standard curve in order to estimate SOD activity in unknown samples.

#### Splenocyte cultures

Spleens of cross-sectional animals were removed, disaggregated, and splenocytes cultured under cell culture conditions equivalent to those used in [42]. Unstimulated splenocytes, taken from 84 cross-sectional animals collected between July and October 2015, were initially used to assay expression by RNASeq (see below). We exposed splenocytes from the remaining cross-sectional animals to stimulation with anti-CD3 antibodies (Hamster Anti-Mouse CD3e, Clone 500A2 from BD Pharmingen, San Diego, California) and anti-CD28 antibodies (Hamster Anti-Mouse CD28, Clone 37.51 from Tombo Biosciences, Kobe, Japan) at concentrations of 2 μg/ml and of 1 μg/ml respectively for 24 hr. Costimulation with anti-CD3 and anti-CD28 antibodies was used to selectively promote the proliferation of T cells [43,44]. We assumed that this would reflect the potential response of T-cell populations *in vivo*. Stimulated splenocytes were used to assay expression by Q-PCR.

#### RNASeq

Full details of the methods used for RNA preparation and sequencing can be found in [3]. Briefly, samples were sequenced on an Illumina HiSeq4000 platform. High-quality reads were mapped against a draft genome for *M. agrestis* (GenBank Accession no. LIQJ00000000) and counted using featureCounts [45]. Genes with average log counts per million, across all samples, of less than one were considered unexpressed and removed from the data (*n* = 8,410). Following filtering, library sizes were recalculated, data were normalized and MDS plots were generated to check for any unusual patterns in the data. The mean library size was 19 million paired-end reads (range = 3 – 71 million paired-end reads).

#### Q-PCR

We used SYBR green based Q-PCR to measure the expression levels of a panel of 18 genes (Appendix 1—table 16) in splenocytes stimulated with anti-CD3 and anti-CD28 antibodies (T cell stimulators) from our cross-sectional animals. We did this, in part, to validate our RNASeq results in an independent dataset. We used the observed expression profile as a general measure of the potential responsiveness of the immune system to an inflammatory stimulation *in vivo*. Although *Fcer1a* is not expressed by T-cells themselves, polymorphism in this gene could be acting indirectly on T-cells through various pathways, including via cytokine signalling, following expression of *Fcer1a* by other cells. The choice of our panel of genes was informed by (i) known immune-associated functions in mice, combined with (ii) significant sensitivity to environmental or intrinsic host variables in our previous studies [9,42] or in a recent DGE analysis of RNASeq data (not reported here), and (iii) the aim of limited redundancy, with each gene representing a different immune pathway.

Primers (20 sets, including 2 endogenous control genes) were designed *de novo* and supplied by PrimerDesign (16 sets) or designed *de novo* in-house (4 sets) and validated (to confirm specific amplification and 100 ± 10% PCR efficiency under assay conditions). All PrimerDesign primer sets were validated under our assay conditions before use. The endogenous control genes (*Ywhaz* and *Sdha*) were selected as a stable pairing from our previous stability analysis of candidate control genes in *M. agrestis* splenocytes [42]. We extracted RNA from splenocytes conserved in RNAlater using the RNAqueous Micro Total RNA Isolation Kit (ThermoFisher), following the manufacturer’s instructions. RNA extracts were DNAse treated and converted to cDNA using the High-Capacity RNA-to-cDNA™ Kit (ThermoFisher), according to manufacturer’s instructions, including reverse transcription negative (RT-) controls for a subsample. SYBR green-based assays were pipetted onto 384 well plates by a robot (Pipetmax, Gilson) using a custom programme and run on a QuantStudio 6-flex Real-Time PCR System (ThermoFisher) at the machine manufacturers default real-time PCR cycling conditions. Reaction size was 10 μl, incorporating 1 μl of template and PrecisionFAST qPCR Master Mix with low ROX and SYBR green (PrimerDesign) and primers at the machine manufacturer’s recommended concentrations. We used two standard plate layouts for assaying, each of which contained a fixed set of target gene expression assays and the two endogenous control gene assays (the same sets of animals being assayed on matched pairs of the standard plate layouts). Unknown samples were assayed in duplicate wells and calibrator samples in triplicate wells and no template controls for each gene were included on each plate. Template cDNA (see above) was diluted 1/20 prior to assay. The calibrator sample (identical on each plate) was created by pooling cDNA derived from across all splenocyte samples. Samples from different sampling groups were dispersed across plate pairs, avoiding confounding of plate with the sampling structure. Gene relative expression values used in analyses are RQ values calculated by the QuantStudio 6-flex machine software according to the ΔΔCt method, indexed to the calibrator sample. Melting curves and amplification plots were individually inspected for each well replicate to confirm specific amplification.

### Longitudinal and cross-sectional data

#### Genotyping

We genotyped both cross-sectional and longitudinal animals for 346 single nucleotide polymorphisms (SNPs) in 127 genes. See [3] for details of the approach used to select these SNPs. Our list included two synonymous and tightly linked (r^2^ = 0.50; D’ = 0.70) SNPs in the gene *Fcer1a* (the alpha chain of the high-affinity receptor for IgE) on scaffold 582 (CADCXT010006977 in ENA accession GCA_902806775; see Appendix 1—table 1 for genomic location information) which we had previously identified as a candidate tolerance gene in a natural population of *M. agrestis*. In this previous work, we had identified four haplotypes at this locus present in our population: GC, AC, AT and GT, at frequencies of 0.12, 0.76, 0.07 and 0.05 respectively. We had also identified the GC haplotype as being of particular interest, given its significantly lower expression level of the transcription factor GATA3 (a biomarker of tolerance to macroparasites in our population) compared to the other haplotypes [3]. We concluded that this haplotype tagged a causal mutation in the coding sequence, or in the up or downstream regulatory regions. DNA was extracted from a tail sample (longitudinal component) or an ear sample (cross-sectional component) taken from the animal using DNeasy Blood and Tissue Kit (Qiagen). Genotyping was then performed by LGC Biosearch Technologies (Hoddesdon, UK; http://www.biosearchtech.com) using the KASP SNP genotyping system. This included negative controls (water) and duplicate samples for validation purposes. The resulting SNP dataset was used for two purposes (i) genotyping individuals within the locus of interest, and (ii) pedigree reconstruction (see below).

#### Pedigree reconstruction

We used a subset of our SNP dataset to reconstruct a pedigree for both cross-sectional and longitudinal animals using the R package *Sequoia* [46]. SNPs which violated the assumptions of Hardy-Weinberg equilibrium were removed from the dataset. For pairs of SNPs in high linkage disequilibrium (most commonly within the same gene), the SNP with the highest minor allele frequency (MAF) was chosen. A minimum MAF cut-off of 0.1 and call rate of > 0.7 was then applied, and any samples for which more than 50% of SNPs were missing were removed. This resulted in a final dataset including 114 SNPs - a sufficient number for very good performance of parentage assignment [46].

Life history information, namely sex and month of birth, was inputted into *Sequoia* where possible. Juvenile voles weighing less than 12 g on first capture were assigned a birth date 2 weeks prior to capture. Juvenile voles weighing between 12 and 15 g on first capture were assigned a birth date 4 weeks prior to capture. Finally, adult voles breeding on first capture, were assigned a birth date 6 weeks prior to capture (minimum age at first breeding) [47,48]. Adult voles not breeding on first capture could not be assigned a birth date, as it was not known whether they had previously bred or not. Virtually all samples (99%) were assigned a sex, and approximately half (54%) were assigned a birth month. As we sampled individuals from across seven different clear-cut areas of the forest, each several kilometres apart, these were assumed to be independent, closed populations with negligible dispersal. Site-specific pedigrees were therefore generated.

We assessed the accuracy of our reconstructed pedigrees by checking whether predicted parent-offspring pairs met expectations given the biology of *M. agrestis*. As expected, the majority of predicted parent-offspring pairs (87%) were born in the same year. We also expected parents and offspring to overlap in space. Again, as expected, the majority of predicted parent-offspring pairs (92%) were, at some point, found along the same transects (horizontal or vertical). We also inspected log10 likelihood ratios (LLRs) for parent pairs as recommended in the user manual for *Sequoia*. Almost all LLRs were positive (*n* = 698/720 or 97% of LLRs) indicating confidence in our assignments. Individuals with vs. without our haplotype of interest did not differ in their probability of appearing in a pedigree (*χ*^2^ = 0.09, d.f. = 1, *p* = 0.76). For each individual that ended up in a pedigree i.e. with one or more relatives recorded (*n* = 652; Site COL = 3; Site BLB = 125; Site GRD = 204; Site CHE = 137; Site RAV = 16; Site SCP = 90; Site HAM = 77), the number of offspring was counted to provide a measure of their reproductive success. Few individuals were first trapped as juveniles, with the majority trapped as adults which had already recruited into the population. Our measure of reproductive success then, more closely resembles the number of recruited (rather than newborn) offspring per individual. Half of individuals present in our pedigrees (*n* = 325) were found to have no offspring. We expect the majority of these to be true zeros (representing actual reproductive failure) as we generally sampled a large proportion of the total population within clear-cuts. We also minimised the chance of false zeros by excluding those individuals (e.g. at the periphery of a study grid) which did not end up in a pedigree because we identified no relatives (including offspring), likely because we had not sampled in the right place.

### Statistical analyses

Not all individuals appeared in all datasets, therefore sample sizes (reported in Table 1) vary between analyses. All analyses were performed in R statistical software version 3.5.2 [49].

**Table 1.**
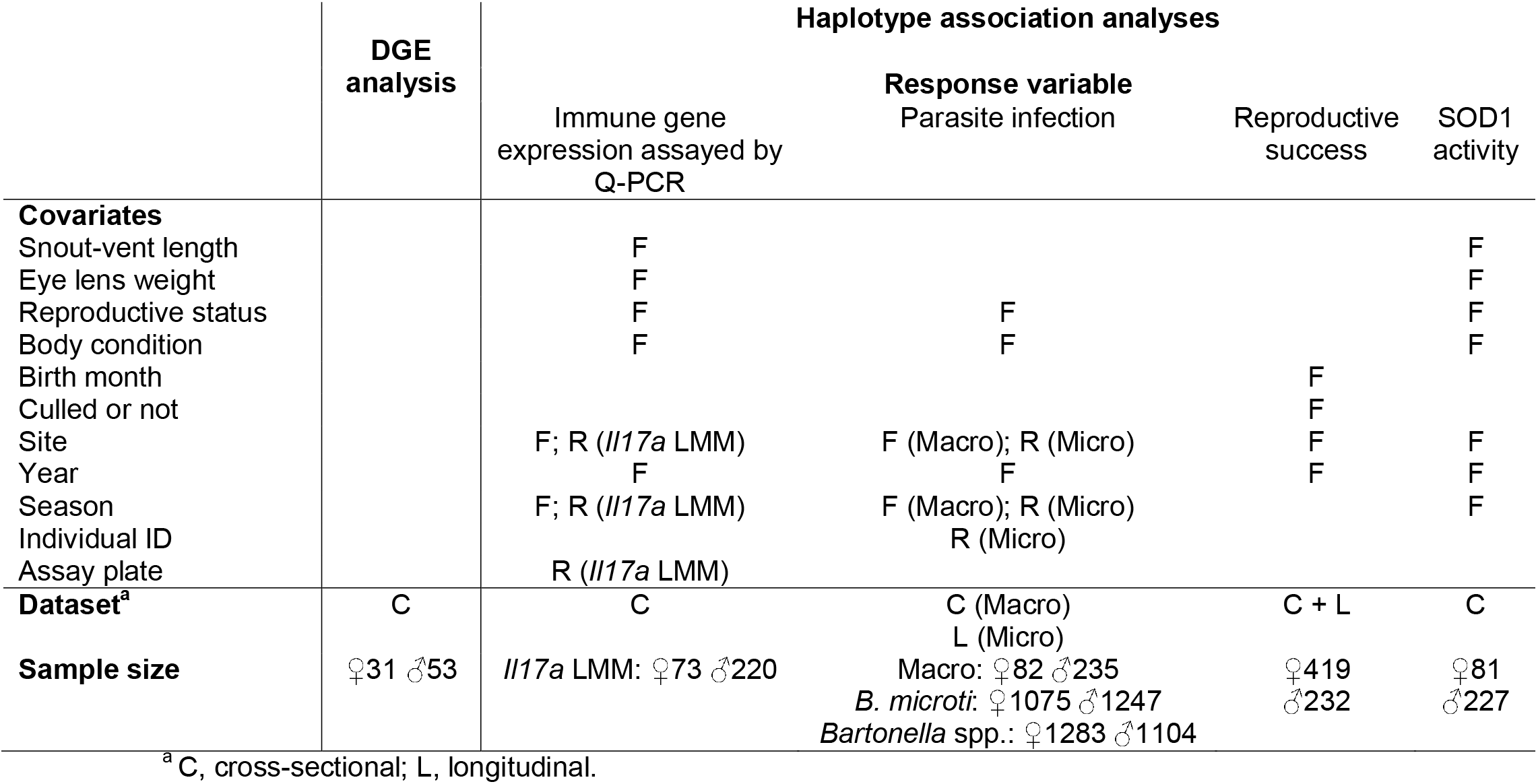
Model specifications including, for each main model: covariates included in the full model, datasets used and sample sizes (F = included as a fixed effect; R = included as a random effect).

#### Differential gene expression analysis

DGE analysis was performed on filtered and normalized count data using the R package edgeR [50], the aim being to identify individual genes which were differentially expressed between those individuals with and without a copy of the GC haplotype (i.e. a dominant model). Only those individuals for which haplotype could be inferred with certainty could be included (*n* = 53 males and *n* = 31 females; none of which were known to have two copies of the GC haplotype, hence the choice of a dominant model). Samples from different sexes were analysed together. To test whether the top-responding genes differed between the sexes, we included a dummy variable with four levels (males with haplotype, males without haplotype, females with haplotype, females without haplotypes) and inspected the contrasts of interest (males with versus without haplotypes; females with versus without haplotype). As this was an exploratory analysis, used in conjunction with more targeted measurements of immune gene expression (see below), model specification was kept as simple as possible and no covariates were initially included. However, in a follow-up DGE analysis we controlled for a key parasite variable (cestode burden).

Having confirmed a sex-dependent effect of the GC haplotype on the expression of some inflammatory genes (see Results) in the initial DGE analysis, we ran separate DGE analyses for males and females and tested more broadly, for enrichment of pro- and anti-inflammatory genes in our results; more specifically, the gene ontology terms, ‘positive regulation of inflammatory response’ (GO:0050729; *n* = 143) and ‘negative regulation of inflammatory response’ (GO:0050728; *n* = 149). The aim here was to answer the question: are pro- or anti-inflammatory genes more highly ranked relative to other genes, when we compare individuals with and without the GC haplotype, and does this also differ between the sexes? This GSEA was performed using the R package limma [51], and genes were ranked on log fold change.

#### Haplotype association analyses

Following the exploratory DGE analysis, the GC haplotype was tested for associations with (i) gene expression assayed by Q-PCR to validate these results, (ii) macro- and microparasite infection to test whether the GC haplotype predicted an individual’s susceptibility to infection, and (iii) reproductive success to test whether the GC haplotype was associated with any reproductive costs. Given the well-established link between inflammation and oxidative stress [52,53], we also tested for an association between the GC haplotype and SOD1 activity.

For all analyses, this was initially attempted using the R package hapassoc [54] in order to maximise sample size (see below). Hapassoc infers haplotypes on the basis of data from single SNPs, and allows likelihood inference of trait associations with resulting SNP haplotypes and other attributes. It adopts a generalized linear model framework and estimates parameters using an expectation–maximization algorithm. Hapassoc models assumed an additive genetic model. If the haplotype combination of an individual cannot, with certainty, be inferred from its genotyping data (i) because it is heterozygous at two or more markers or (ii) because it has missing data for a single marker, the approach implemented in hapassoc is to consider all possible haplotype combinations for that individual. Standard errors accounting for this added uncertainty are calculated using the Louis’ method [55]. We compared the GC haplotype against the other two major haplotypes (AC and AT). Another haplotype, the GT haplotype, was identified in the population but this was present at such low frequencies (frequency ≤ 0.01 among individuals for which haplotype could be inferred with certainty) that it was omitted from all analyses. Results reported in the text for macroparasites and SOD1 activity come from these hapassoc models.

However, there are some restrictions on model specification within hapassoc (e.g. random terms cannot be included, limited choice of error distributions), so this was followed up with regression models for some analyses. As in the DGE analysis, these regression models only included those individuals whose haplotype combination could be inferred with certainty. Results reported in the text for gene expression assayed by Q-PCR, microparasites and reproductive success come from these regression models. Again, as in the DGE analysis, genotype was coded as the presence or absence of the GC haplotype (i.e. a dominant model). Regression models were run using the R package lme4 [56] or glmmADMB [57,58].

Table 1 provides a summary of (hapassoc or regression) model specifications. As we found evidence for a sex-dependent effect in the initial DGE analysis, with the exception of the model for reproductive success (see below), all models included a genotype by sex interaction which was retained if it improved model fit. All models also accounted for other biological and technical covariates (the choice of which was informed by our previous work, the literature and/or our experimental design). Full regression models (including all covariates and interactions of interest) were reduced using backward stepwise deletion of non-significant terms to minimize Akaike’s information criterion (AIC), following the drop1 function in the lme4 package. This was not possible for hapassoc models. Summary tables including all estimates, standard errors and *z*-statistics are included in Appendix 1. Associated significance values are also included in these tables, except for LMMs and GLMMs, for which significance values were based on likelihood ratio tests (LRTs) and are reported in the main text. Throughout, residuals from regression models were checked for approximate normality and homoscedasticity, and all covariates were tested for independence using variance inflation factors (all VIFs < 3).

#### Association between GC haplotype and immune gene expression assayed by Q-PCR

As we ran a total of 18 hapassoc models (one model per gene), the Benjamini and Hochberg method of correction was applied to all *p*-values [59]. Resulting *q*-values (FDR-corrected *p*-values) are reported, alongside original *p*-values (Appendix 1—table 6). A hapassoc model including a genotype by sex interaction was tested against a null model without an interaction term. A Gamma error distribution and log link were used. Other covariates considered potential drivers of immune gene expression and included in both models, were informed by our previous work [42]. These included snout-vent length (SVL), eye lens weight (categorised into seven intervals; SVL and eye lens weight capture the combined influence of age and historical growth trajectory), reproductive status (males were considered to be reproductively active if they had descended testes; females if they were pregnant or had perforate vaginas) and body condition (estimated by regressing body weight against life history stage, SVL and its quadratic term). Site, year and season (four levels, designated as spring [March & April], early summer [May and June], late summer [July and August] and autumn [September and October]) were included to account for any spatial and/or temporal autocorrelation. All covariates were included as fixed effects. We did not use a multidimensional approach (such as principal component analysis) because of limited redundancy in our panel of genes.

In order to confirm these results, a linear mixed effects model (LMM) was run for a single immune gene for which expression appeared to be associated with genotype (*q* = 0.037). The LMM included random terms for site and season, as well as assay plate number (Table 1). The latter was included to account for nonindependence due to immunological assaying structure. All other covariates were the same as the hapassoc model. A Yeo-Johnson transformation (with λ = −2) was used to achieve more normal and homoscedastic residuals [60].

#### Association between GC haplotype and parasite infection

The three macroparasite measurements taken from cross-sectional animals (counts of ticks, fleas and cestodes) were log-transformed (log_10_ [*x* +1]) and summarised as a single principal component (explaining 39% of total variation; Appendix 1—table 17) to avoid difficulties in interpretation due to multiple testing. See [3] and [9] for full details of this approach. This combined measure of macroparasite burden was modelled using a hapassoc model with a Gaussian error distribution.

Microparasite infection status was assessed multiple times for the majority of individuals in the longitudinal component of the study (mean = 2.8; range = 1-11). Due to these repeated measures, microparasite infection could not be modelled using hapassoc. Instead, to test for an association between the GC haplotype and the probability of an individual being infected with a microparasite, we ran a GLMM with a binary response (infected or not), log link and a random term for individual. Other covariates, considered potential drivers of both macro and microparasite infection were, again, informed by our previous work [3,9,61]. These included body condition, reproductive status, year, season and site (Table 1). Season and site were included as random terms in the GLMM for microparasites, as was individual identity, to account for nonindependence due to repeat sampling.

#### Association between GC haplotype and reproductive success

Our measure of reproductive success was zero-inflated (50% zeros). This is consistent with a previous study of the closely-related common vole (*Microtus arvalis*) [62]. A Poisson error distribution was therefore deemed inappropriate and it could not be modelled using hapassoc. Instead, to test for an association between the GC haplotype and reproductive success, we ran a GLM with a quasi-Poisson error distribution. This distribution has been previously used to model predictors of reproductive success in other organisms, and accounts for the overdispersion caused by excess zeros.

We only recorded a reproductive failure (i.e. zero reproductive success) for those individuals with some other familial relationship in our pedigree, purposefully omitting those individuals (e.g. at the periphery of a study grid) for which we may have recorded no relatives (including offspring) simply because we had not sampled in the right place. Therefore, we expect all (or most) of our zeros to represent actual reproductive failure. For this reason, zero-inflated models were deemed inappropriate.

As detailed in the Results, we tried to run a single quasi-Poisson GLM with both sexes but resulting residual variances differed significantly between the sexes (F test to compare variances of two samples; *p* = 0.02), making it impossible to formally test for a genotype by sex interaction. Instead, we ran a separate model for each sex. We included birth month as a covariate in this model, given that autumn-born voles (of the closely-related common vole) have been shown to have a lower chance to reproduce than spring-born voles [62]. Other covariates included in this model were, whether or not an individual was culled for the cross-sectional component of this study (again, reducing the opportunity to reproduce), site and the year in which an individual was most frequently captured (Table 1). In a follow-up analysis, we also controlled for *Babesia microti* and *Bartonella* spp. infection status. More specifically, we included the proportion of samples taken from an individual that were *Babesia*-positive, and the proportion of samples taken from an individual that were *Bartonella-positive*. All covariates were included as fixed effects. Only a single female was trapped in one of the sites (COL) and consequently caused convergence problems in the female model. This female was therefore omitted.

#### Association between GC haplotype and SOD1 activity

SOD1 activity was modelled using a hapassoc model with a Gaussian error distribution. Other covariates, considered potential drivers of SOD1 activity, were informed by the literature. Previous studies on wild rodents have shown that antioxidant levels increase with both reproductive effort [63] and with age [64]. Studies on birds have also shown that improvements in body condition are often accompanied by increases in antioxidant activity, for example in response to supplemental feeding [65]. We therefore included SVL, eye lens weight, reproductive status and body condition as covariates in our model. As in other models, we included site, year and season to account for spatial and/or temporal autocorrelation in our data (Table 1). All covariates were included as fixed effects.

## Acknowledgements

We would like to thank all those involved in obtaining and processing samples from the field: Rebecca Turner, Lukasz Lukomski, Stephen Price, Sarah Gore, Ed Parker, Maria Capstick, Noelia Dominguez Alvarez, Susan Withenshaw, William Foster, Ann Lowe and Benoit Poulin. We would also like to thank the Forestry Commission for access to the study sites and the Centre for Genomic Research at the University of Liverpool for sequencing samples. This research was funded by the Natural Environment Research Council (NERC) award NE/L013452/1 to S.P., M.B., J.A.J. and J.E.B.

## Data availability

RNASeq data have been deposited in the European Nucleotide Archive (study accession number PRJEB51626). Longitudinal phenotypic data are available from the NERC EDS Environmental Information Data Centre (doi:10.5285/e5854431-6fa4-4ff0-aa02-3de68763c952). Cross-sectional phenotypic data are available from the University of Liverpool’s Research Data Catalogue (doi:10.17638/datacat.liverpool.ac.uk/1850). Genotype data are also available from the University of Liverpool’s Research Data Catalogue (doi:10.17638/datacat.liverpool.ac.uk/1849).

## Appendix 1

**Appendix 1—table 1.** Position of SNPs and other key features in the *Fcer1a* gene. Features lie in scaffold CADCXT010006977 within assembly GCA_902806775.

| Feature | Start | End |
| --- | --- | --- |
| *Fcer1a* gene | 10745528 | 10745528 |
| Exon 5 | 10745528 | 10745881 |
| Exon 4 | 10746818 | 10747076 |
| Exon 3 | 10748609 | 10748864 |
| Exon 2 | 10750044 | 10750065 |
| Exon 1 | 10750465 | 10750642 |
| 5' UTR | 10750520 | 10750642 |
| CDS | 10750465 | 10750520 |
| CDS | 10750044 | 10750065 |
| CDS | 10748609 | 10748864 |
| CDS | 10746818 | 10747076 |
| CDS | 10745729 | 10745881 |
| 3' UTR | 10745528 | 10745729 |
| SNP 2 | 10748718 | 10748719 |
| SNP 1 | 10746846 | 10746847 |

**Appendix 1—table 2.**
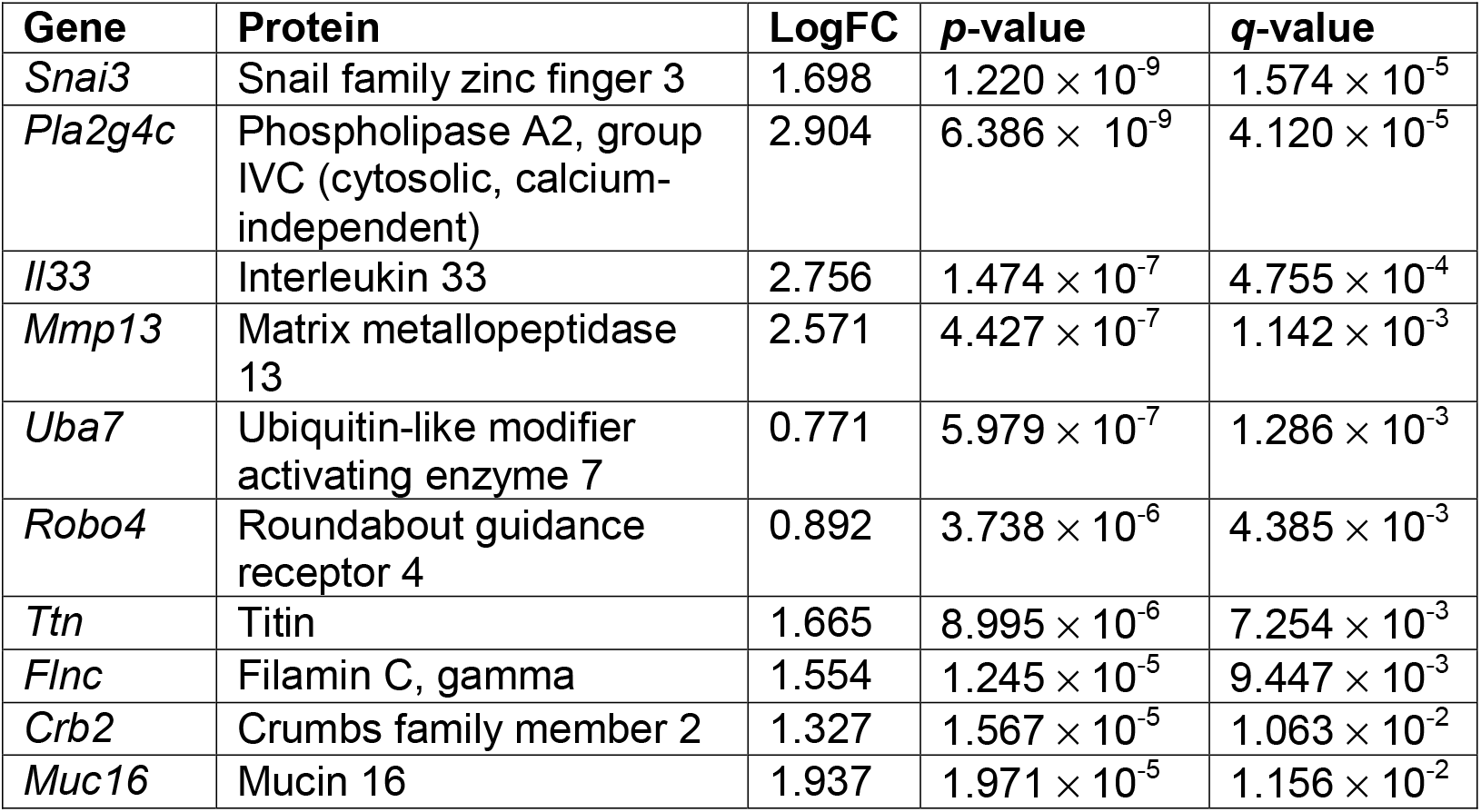
Top 10 annotated genes which were differentially expressed between males with vs. without the GC haplotype, including associated log fold changes (logFC), *p*-values and *q*-values (FDR-corrected *p*-values).

**Appendix 1—table 3.**
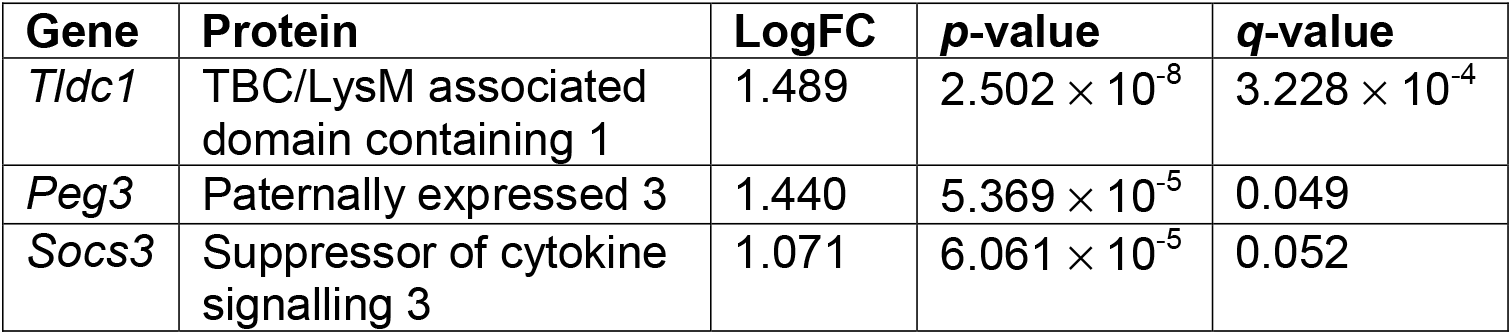
Annotated genes which were differentially expressed (*q* ≤ 0.1) between females with vs. without the GC haplotype, including associated log fold changes (logFC), *p*-values and *q*-values (FDR-corrected *p*-values).

**Appendix 1—table 4.**
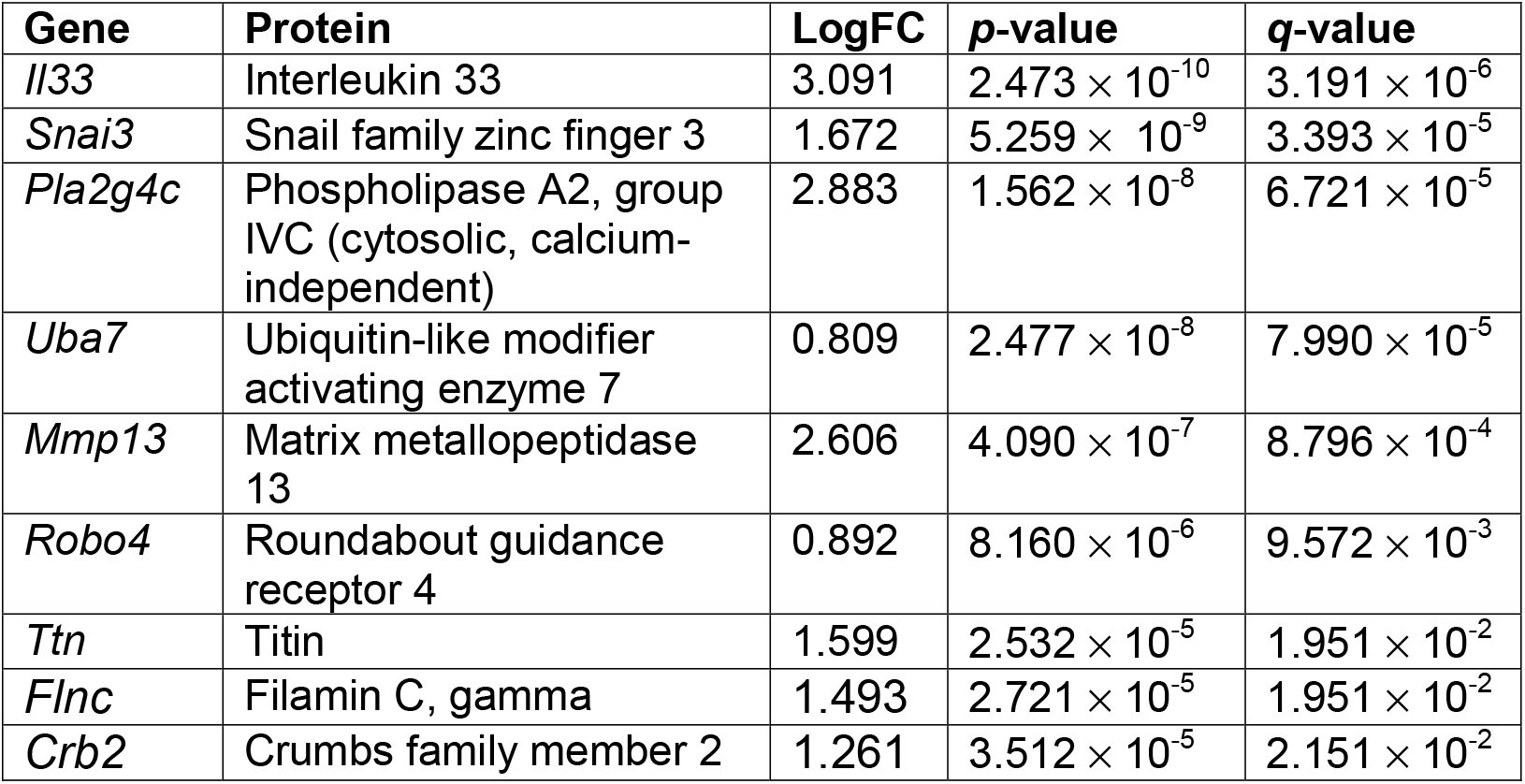
Top 10 annotated genes which were differentially expressed between males with vs. without the GC haplotype when controlling for cestode burden, including associated log fold changes (logFC), *p*-values and *q*-values (FDR-corrected *p*-values).

**Appendix 1—table 5.**
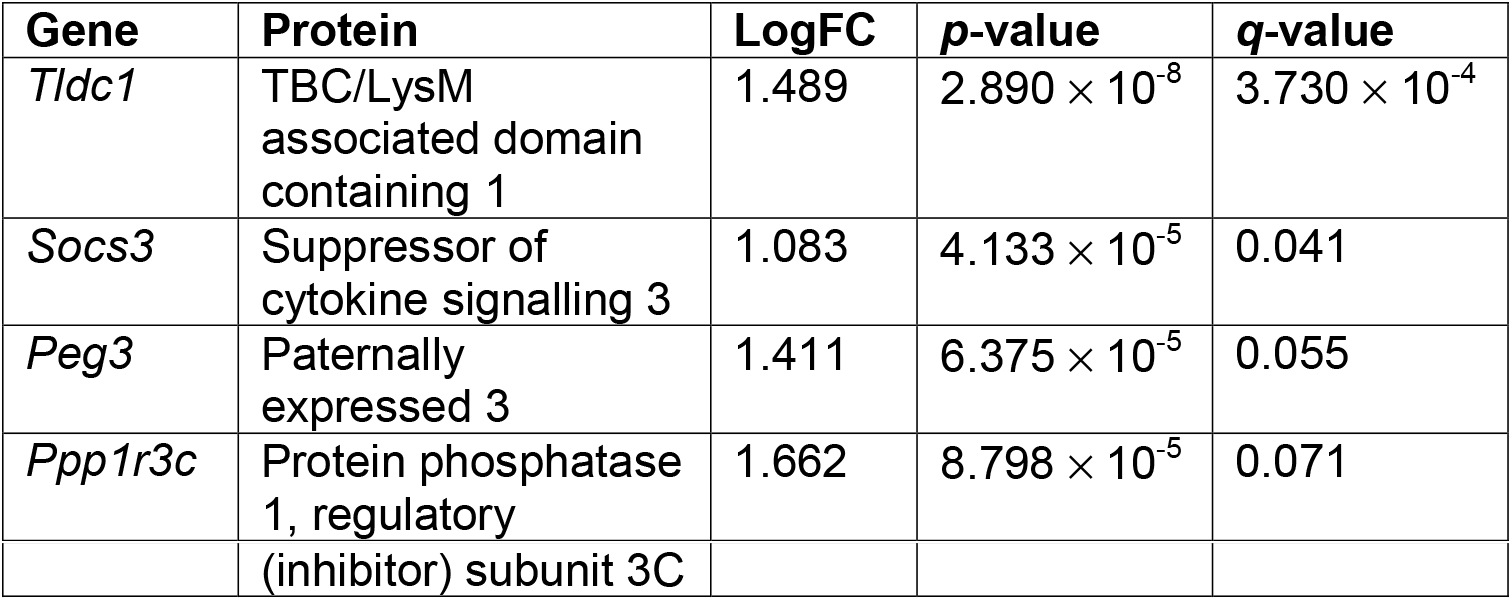
Annotated genes which were differentially expressed (*q* ≤ 0.1) between females with vs. without the GC haplotype when controlling for cestode burden, including associated log fold changes (logFC), *p*-values and *q*-values (FDR-corrected *p*-values).

**Appendix 1—table 6.** Significance values from hapassoc models for expression of 18 genes (assayed by Q-PCR) in splenocytes. Splenocytes were stimulated with anti-CD3 and anti-CD28 antibodies in order to promote the proliferation of T cells. *q*-values (FDR-corrected *p*-values) are reported alongside original *p*-values for the genotype by sex interaction.

| Gene | *p*-value | *q*-value |
| --- | --- | --- |
| *Cd4* | 0.124 | 0.822 |
| *Cd8a* | 0.744 | 0.866 |
| *Foxp3* | 0.499 | 0.845 |
| *Gata3* | 0.563 | 0.845 |
| *Il10* | 0.866 | 0.866 |
| *Mpo* | 0.173 | 0.822 |
| *Tbx21* | 0.650 | 0.866 |
| *Tgfb1* | 0.271 | 0.822 |
| *Ifng* | 0.399 | 0.822 |
| *Il17a* | 0.002 | 0.037 |
| *Il1b* | 0.749 | 0.866 |
| *Il6* | 0.282 | 0.822 |
| *Ms4a1* | 0.281 | 0.822 |
| *Orai1* | 0.353 | 0.822 |
| *Tnfa* | 0.858 | 0.866 |
| *Il2* | 0.411 | 0.822 |
| *Apobr* | 0.857 | 0.866 |
| *Arg1* | 0.552 | 0.845 |

**Appendix 1—table 7.** Estimates, standard errors and *z*-statistics from best LMM for Yeo-Johnson-transformed *Il17a* expression levels.

|  | Estimate | SE | z |
| --- | --- | --- | --- |
| (Intercept) | 0.107 | 0.023 | 4.61 |
| Genotype | -0.115 | 0.037 | -3.10 |
| Sex Male | -0.055 | 0.019 | -2.89 |
| Year 2016 | 0.084 | 0.018 | 4.60 |
| Genotype x Sex Male | 0.143 | 0.044 | 3.25 |

**Appendix 1—table 8.** Effect sizes, standard errors, *z*-statistics and associated significance from Gaussian hapassoc model for SOD1 activity.

|  | Estimate | SE | z | *p*-value |
| --- | --- | --- | --- | --- |
| (Intercept) | 0.614 | 1.514 | 0.406 | 0.685 |
| hAT | -0.453 | 0.223 | -2.032 | 0.042 |
| Reproductive status Active | -0.112 | 0.266 | -0.422 | 0.673 |
| Sex Male | 0.031 | 0.250 | 0.124 | 0.901 |
| SVL | 0.016 | 0.017 | 0.945 | 0.345 |
| Lens weight 2 | 0.568 | 0.531 | 1.070 | 0.284 |
| Lens weight 3 | 0.440 | 0.549 | 0.801 | 0.423 |
| Lens weight 4 | 0.445 | 0.587 | 0.758 | 0.448 |
| Lens weight 5 | 0.415 | 0.653 | 0.636 | 0.525 |
| Lens weight 6 | 0.301 | 0.752 | 0.400 | 0.689 |
| Lens weight 7 | -0.307 | 1.095 | -0.280 | 0.779 |
| Body condition | 0.020 | 0.030 | 0.677 | 0.498 |
| Site CHE | 1.429 | 0.431 | 3.316 | 0.001 |
| Site COL | 2.675 | 1.099 | 2.434 | 0.015 |
| Site GRD | 1.476 | 0.437 | 3.379 | 0.001 |
| Site RAV | 0.904 | 0.918 | 0.984 | 0.325 |
| Site SCP | 1.107 | 0.419 | 2.643 | 0.008 |
| Year 2016 | 0.593 | 0.242 | 2.457 | 0.014 |
| Season Early Summer | -0.392 | 0.389 | -1.007 | 0.314 |
| Season Late Summer | -0.098 | 0.266 | -0.369 | 0.712 |
| Season Spring | -0.969 | 0.383 | -2.529 | 0.011 |

**Appendix 1—table 9.** Effect sizes, standard errors and *z*-statistics from best Binomial GLMM for probability of infection with *Babesia microti*.

|  | Estimate | SE | z |
| --- | --- | --- | --- |
| (Intercept) | -2.214 | 0.377 | -5.87 |
| Genotype | 1.254 | 0.416 | 3.02 |
| Sex Male | 0.957 | 0.229 | 4.19 |
| Year 2016 | 1.260 | 0.277 | 4.55 |
| Year 2017 | 1.047 | 0.302 | 3.46 |
| Reproductive status Active | 0.754 | 0.144 | 5.24 |
| Body condition | 0.044 | 0.016 | 2.76 |
| Genotype x Sex Male | -1.116 | 0.615 | -1.82 |

**Appendix 1—table 10.** Effect sizes, standard errors and *z*-statistics from best Binomial GLMM for probability of infection with *Bartonella* spp.

|  | Estimate | SE | z |
| --- | --- | --- | --- |
| (Intercept) | 0.236 | 0.369 | 0.64 |
| Year 2016 | 0.740 | 0.135 | 5.49 |
| Year 2017 | 2.280 | 0.175 | 13.01 |

**Appendix 1—table 11.** Effect sizes, standard errors, *z*-statistics and associated significance from Gaussian hapassoc for macroparasite infection summarised by a single principal component.

|  | Estimate | SE | z | p-value |
| --- | --- | --- | --- | --- |
| (Intercept) | 0.050 | 0.331 | 0.152 | 0.879 |
| hAC | -0.102 | 0.130 | -0.786 | 0.432 |
| hAT | -0.108 | 0.158 | -0.682 | 0.495 |
| Sex Male | -0.271 | 0.126 | -2.146 | 0.032 |
| Season Early Summer | -0.103 | 0.182 | -0.567 | 0.571 |
| Season Late Summer | 0.012 | 0.132 | 0.090 | 0.929 |
| Season Spring | 0.327 | 0.176 | 1.854 | 0.064 |
| Reproductive status Active | -0.222 | 0.119 | -1.868 | 0.062 |
| Body condition | -0.019 | 0.014 | -1.328 | 0.184 |
| Year 2016 | -0.473 | 0.124 | -3.810 | 0.000 |
| Site CHE | 0.714 | 0.226 | 3.157 | 0.002 |
| Site COL | 0.762 | 0.575 | 1.327 | 0.185 |
| Site GRD | 0.707 | 0.229 | 3.090 | 0.002 |
| Site RAV | 1.100 | 0.481 | 2.288 | 0.022 |
| Site SCP | 0.747 | 0.220 | 3.389 | 0.001 |

**Appendix 1—table 12.** Effect sizes, standard errors, *z*-statistics and associated significance from best quasi-Poisson GLM for reproductive success in males.

|  | Estimate | SE | z | p-value |
| --- | --- | --- | --- | --- |
| (Intercept) | 1.743 | 0.243 | 7.171 | $1.05 \times 10^{-11}$ |
| Genotype | -0.484 | 0.237 | -2.042 | 0.042 |
| Year 2016 | 0.592 | 0.185 | 3.193 | 0.002 |
| Year 2017 | -0.437 | 0.199 | -2.193 | 0.029 |
| Birth month | -0.358 | 0.048 | -7.449 | $1.97 \times 10^{-12}$ |
| Culled Yes | 0.422 | 0.150 | 2.809 | 0.005 |

**Appendix 1—table 13.**
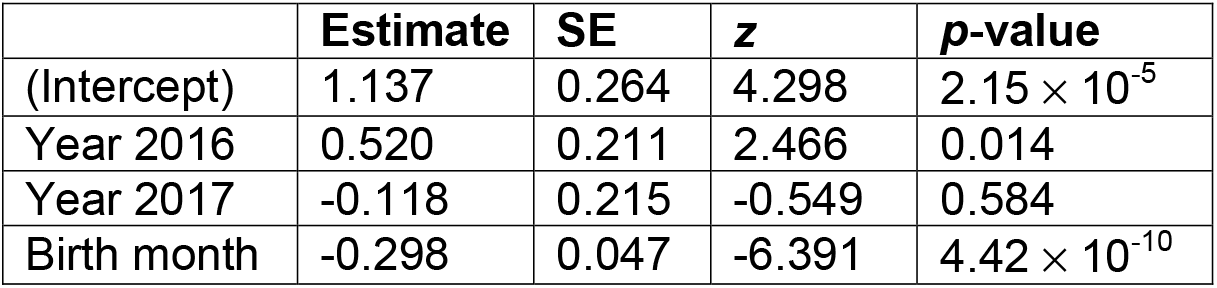
Effect sizes, standard errors, *z*-statistics and associated significance from best quasi-Poisson GLM for reproductive success in females.

**Appendix 1—table 14.** Effect sizes, standard errors, *z*-statistics and associated significance from best quasi-Poisson GLM for reproductive success in males, when controlling for *Babesia microti* and *Bartonella* spp. infection.

|  | Estimate | SE | z | p-value |
| --- | --- | --- | --- | --- |
| (Intercept) | 1.467 | 0.418 | 3.513 | $7.64 \times 10^{-4}$ |
| Genotype | -1.081 | 0.425 | -2.544 | 0.013 |
| Year 2016 | 0.238 | 0.279 | 0.854 | 0.396 |
| Year 2017 | -0.912 | 0.386 | -2.338 | 0.022 |
| Birth month | -0.464 | 0.091 | -5.090 | $2.69 \times 10^{-6}$ |
| Proportion of samples *Babesia*-positive | 0.849 | 0.276 | 3.076 | 0.003 |
| Proportion of samples | 0.997 | 0.440 | 2.268 | 0.026 |
| *Bartonella*-positive |  |  |  |  |

**Appendix 1—table 15.** Effect sizes, standard errors, *z*-statistics and associated significance from best quasi-Poisson GLM for reproductive success in females, when controlling for *Babesia microti* and *Bartonella* spp. infection.

|  | Estimate | SE | z | p-value |
| --- | --- | --- | --- | --- |
| (Intercept) | 1.483 | 0.305 | 4.867 | $3.20 \times 10^{-6}$ |
| Birth month | -0.310 | 0.073 | -4.229 | $4.38 \times 10^{-5}$ |
| Proportion of samples *Babesia*-positive | 0.569 | 0.247 | 2.308 | 0.0226 |

**Appendix 1—table 16.** Panel of 18 genes for which expression levels in splenocytes stimulated with anti-CD3 and anti-CD28 antibodies were measured using two-step reverse transcription quantitative PCR (Q-PCR).

| Gene | Protein |
| --- | --- |
| *Cd4* | T-cell surface glycoprotein CD4 |
| *Cd8a* | T-cell surface glycoprotein CD8 alpha chain |
| *Foxp3* | Forkhead box protein P3 |
| *Gata3* | GATA binding protein 3 |
| *Il10* | Interleukin-10 |
| *Mpo* | Myeloperoxidase |
| *Tbx21* | T-box transcription factor TBX21 |
| *Tgfb1* | Transforming growth factor beta 1 |
| *Ifng* | Interferon gamma |
| *Il17a* | Interleukin-17a |
| *Il1b* | Interleukin-1 beta |
| *Il6* | Interleukin-6 |
| *Ms4a1* | B-lymphocyte antigen CD20 |
| *Orai1* | Calcium release-activated calcium channel protein 1 |
| *Tnfa* | Tumor necrosis factor alpha |
| *Il2* | Interleukin-2 |
| *Apobr* | Apolipoprotein B receptor |
| *Arg1* | Arginase-1 |

**Appendix 1—table 17.**
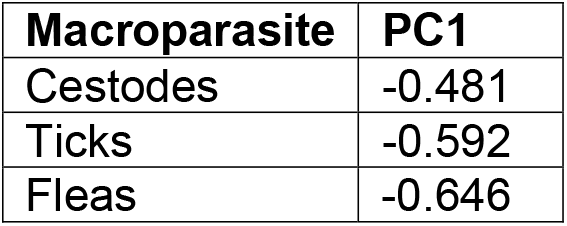
Loadings from principal component analysis summarising infection by macroparasites (ticks, fleas and cestodes)

